# Dispersal behavior in a cold-water coral is orchestrated via stage and species-specific physiology

**DOI:** 10.64898/2026.06.03.729899

**Authors:** Marte Lønnum, Johan Hovland, Madeline M. Schuldt, Eirik Stamland Nilssen, José Davila-Velderrain, Johanna Järnegren, Lena van Giesen

## Abstract

Corals form important ecosystems that serve as habitat for numerous marine species. Being sessile, adult corals are exposed to changing environments without the means to relocate. Species dispersal is therefore restricted to the motile larval lifestage. How do microscopic larvae achieve reliable dispersal and conquest of novel habitats under time pressure and unpredictable environmental conditions? Here we show an unexpected diversity of behaviors in the cold-water coral *Lophelia pertusa*. Anatomical and behavioral changes of coral planula promote a change from neutral, passive buoyancy in the dispersal phase, to active swimming and search behavior during competency. As lipids are metabolized and sensory abilities develop, the coral larvae drastically change their motility patterns. Comparative analysis with a poorly dispersing, lecithotrophic anthozoan larvae reveals that developmentally timed sensory integration is conserved between species, but the behavioral modes and sensory responses are adapted to their particular ecology.

## Introduction

Coral reefs form the foundation for some of the most diverse marine ecosystems on the planet, serving as home to myriads of marine species. While enigmatic, tropical coral reef ecosystems are well studied, the biology of their counterparts in the deep, dark oceans is less well understood^1^. Cold-water corals (CWCs) occur in the deep sea, forming massive, three-dimensional reef structures that are hotspots for biodiversity^1–3^. The most common, framework-forming, stony cold-water coral with a nearly global distribution is *Lophelia pertusa* (syn. *Desmophyllum pertusum*) (**Fig. 1A,B**)^4,5^. Corals are sessile invertebrates bound for life to their habitat, and with its longevity and slow growth, *Lophelia pertusa* is particularly sensitive to environmental changes and physical damage^6^.

**Figure 1.**
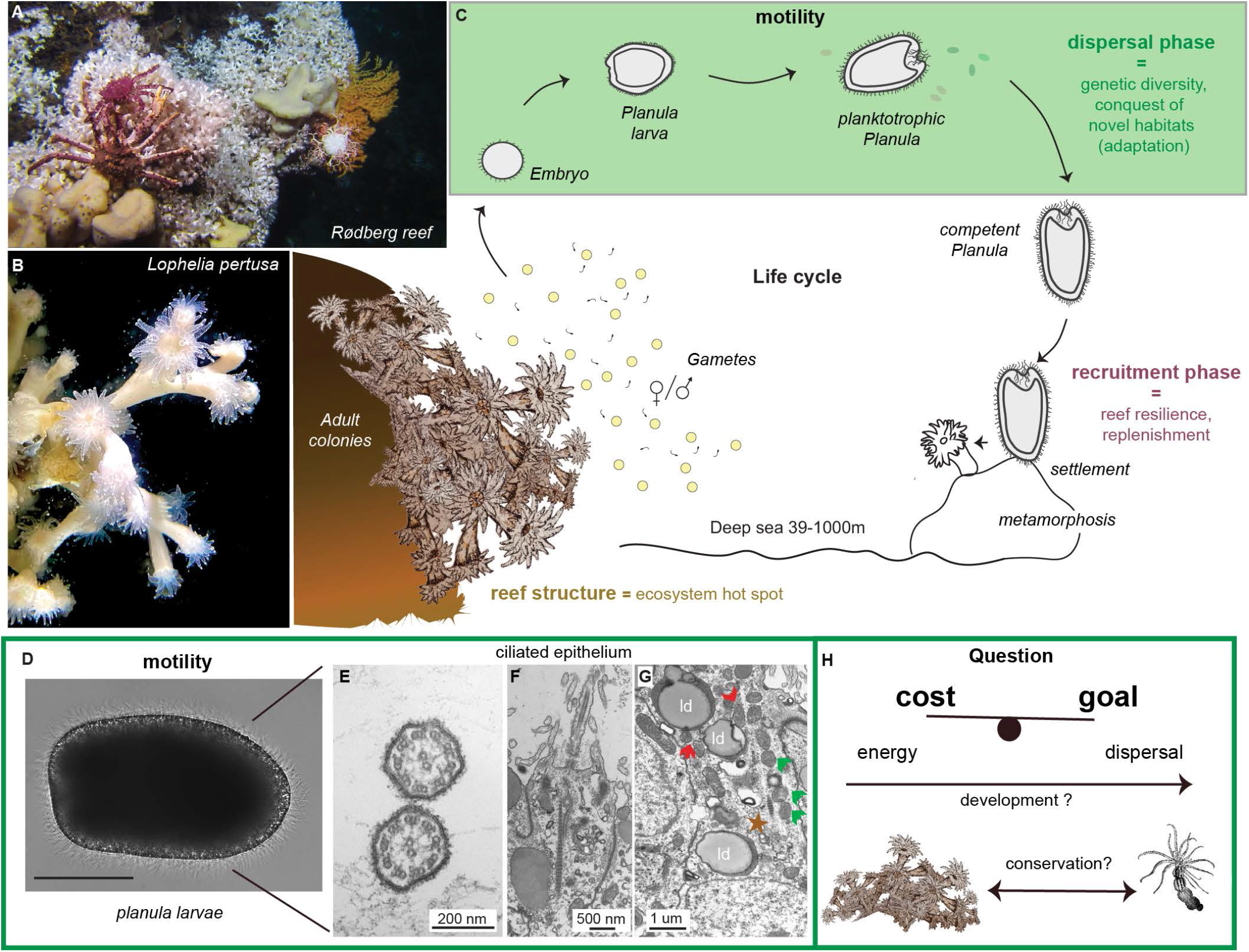
*Lophelia pertusa* dispersal through motile planula. **A)** Coral garden at the Rødberg reef with *Lophelia pertusa* forming reef structures which serve as habitat for a plethora of other species. **B)** Laboratory housed specimens are used for annual spawning events. **C)** Adult colonies release gametes and after successful fertilization and embryos later develop into motile planula larvae. The dispersal phase of otherwise sessile corals is of paramount importance for genetic diversity (reef connectivity). **D)** Planula (31dpf), with densely ciliated epithelium and **E)** cross-section through motile cilia and **F-G)** transversal section through ciliated cells with lipid droplets (ld), peridroplet mitochondria (red arrow), ciliary base (star) and mitochondria by the base (green arrows). **H)** How is dispersal balanced with energetic cost across cnidarian planulae?

Corals have a complex lifecycle, including a motile larval stage, which is critical for replenishment of reefs and conquest of new habitats (**Fig. 1C**). When contemplating coral ecosystem resilience, both retention and dispersal of the larvae are crucial. How do larvae “decide” where to go? Dispersal is critical for genetic diversity and adaptability, but reaching distant locations in the vastness of the ocean exceeds the physical capabilities of microscopic planulae. Relocalization is therefore often determined by water movement and ocean currents. Consequently, developmental length and maternally deposited energy resources play a major role in the distance that can be travelled^7^. Yet, the larva is no passive particle: both settling rates and motility are fluid and physiologically regulated. It follows, therefore, that physiochemical conditions, physiological makeup, and the resulting behavior are determinants of coral larval positioning in the water column, which in turn will affect its dispersal capacity^8–10^.

Knowledge about larval biology is important to understand dispersal potential^11,12^, protect these marine ecosystems^1^, and grasp how deep-sea organisms deal with their particular environmental constraints. However, cold-water corals live in extreme environments and are difficult to access. Previous research has started to investigate *Lophelia pertusa’s* embryonic and larval development^13–15^, but knowledge about how behavioral, physiological, and anatomical adaptations conspire to achieve successful dispersal is currently lacking. How are these different aspects changing over development? Are they adapted to the ecology of the individual species?

Leveraging our rare access to both adult *Lophelia* colonies in the deep sea and laboratory facilities to induce spawning and time development (**Fig. 1A,B**), here we present the first comprehensive data set of coral larval behavior and anatomy over the course of 2.5 months of development. We employ behavioral tracking assays to study the behavior of 5 different developmental stages of the cold-water coral larvae. Developmental transitions in vertical positioning are supported by different motility strategies, as observed when tracking detailed movement of planulae in the horizontal plane. The behavioral modes in turn are products of the animal’s physiology and anatomy which support species-, and stage-specific dispersal. Developmentally timed sensory integration is conserved between species with or without vast dispersal capability, but the sensory responses and behavioral modes seem adapted to their particular ecology.

## Results

### *Lophelia pertusa* as a model for cold-water coral dispersal

How do microscopic planulae achieve reliable dispersal and conquest of novel habitats under time pressure and with unpredictable environmental conditions in a large body of water? Here, we used the larvae of the cosmopolitan cold-water coral (CWC) *Lophelia pertusa* **(Fig. 1)** to understand how successful dispersal can occur despite these challenges.

*Lophelia* planulae larvae are densely ciliated, and motility is achieved through the synchronous beating of these motorized organelles (**Fig. 1D** and **supplementary video 1 and 2**). While the metachronal beating and the positioning of the cilia may be optimized for low energy expenditure^16,17^, viscous forces in seawater make swimming energetically costly for very small organisms^10,18^. The 9+2 microtubule structure found in the cilia (**Fig. 1E,F**) forms the characteristic dynein-based, ATP-dependent motor of these organelles. Accumulations of mitochondria near the base of the cilium (**Fig. 1G** star=ciliary base, green arrows=mitochondria), and peridroplet mitochondria (**Fig. 1G** red arrows) are consistent with the high metabolic demand of these cells. The question then is, how does the animal balance the cost of motility and the goal to disperse (**Fig. 1H**)? Are these aspects solved in the same way over development and conserved across species or are specialized strategies employed?

### Larvae assume different vertical distributions throughout development

Depth distribution of coral larvae is an important factor for the dynamics of dispersal and retention^9,19,20^ and might serve as an indicator for stage-specific dispersal drive (**Fig. 2A**). To determine vertical positioning across development, we designed a custom chamber with restricted depth (z-plane=1cm) and homogeneous illumination to record and track the animal’s behavior in a 10 cm deep water column over several minutes (**Fig. 2B**).

**Figure 2.**
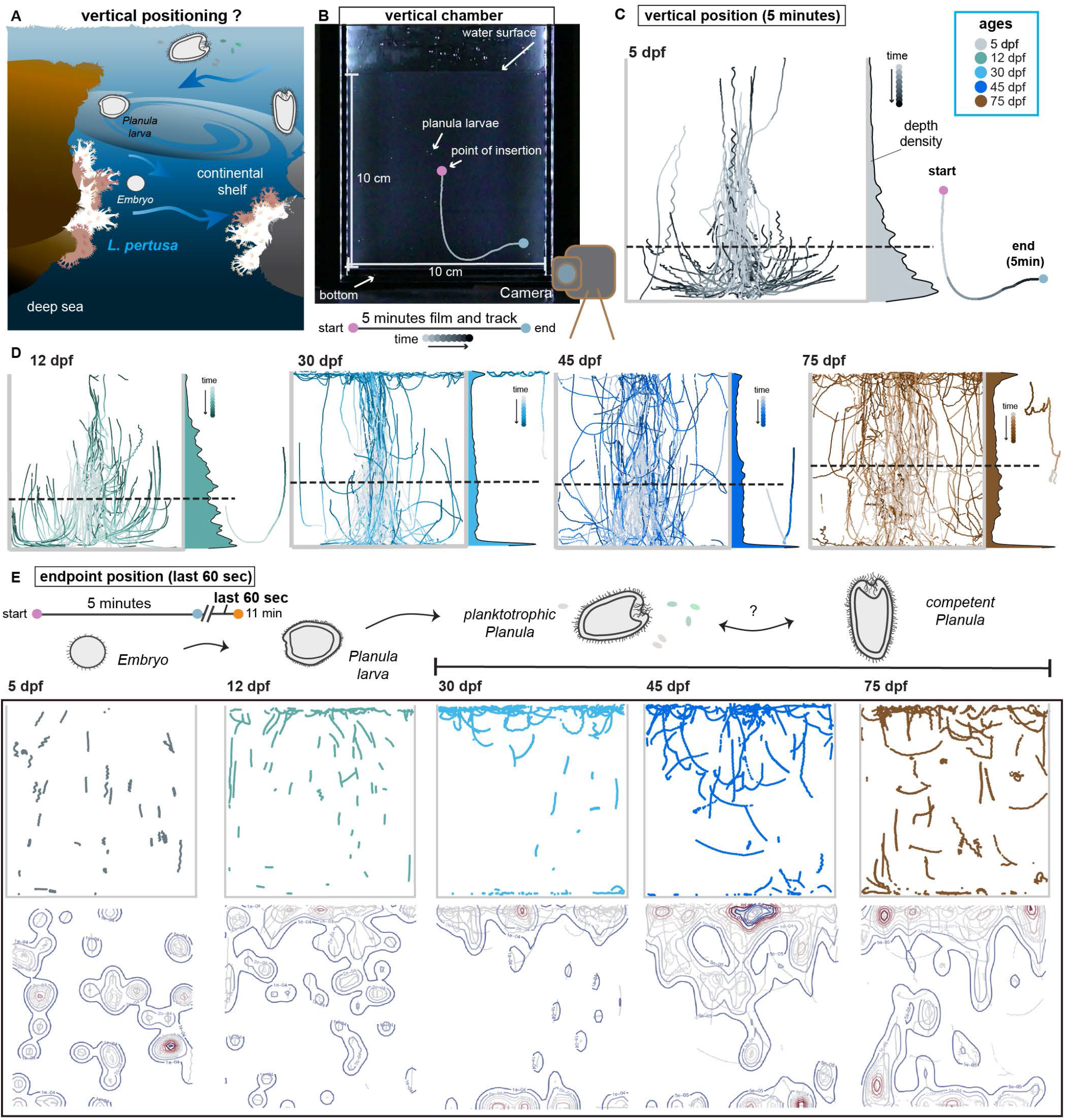
Vertical positioning changes over development. **A)** Natural vertical position is unknown in *Lophelia* planula. **B)** Experimental setup (top) and timeline (bottom) used to track the animals. Planula larvae in suspension and track with insertion point (pink) and end point (blue). **C)** Overlaid tracks color coded in 30-second increments for 5dpf (16N, 89n). The vertical position is shown for five minutes after insertion into the chamber. Density plots, with dotted lines at the mean, show the larval depth distribution during these five minutes (5dpf: 7.89cm). Legend denotes time coloration, individual track on the right-hand representative. **D**) Plots for all larvae aged 12dpf (16N, 99n), 30dpf (18N, 113n), 45dpf (17N,106n) and 75dpf (15N, 105n) (Average depth per age (cm): 12dpf: 7.04, 30dpf: 6.36, 45dpf: 6.32, 75dpf: 5.28). **E)** Position (top) and density distribution (bottom) of the same animals for the last minute (10-11th minute). See also **Table 2**.

Indeed, larval position in the water column is drastically changing for the five chosen ages (5-, 12-, 30-, 45- and 75 days post fertilization(dpf)) (**Fig. 2C,D; Supplementary Fig. S1A-C**, with KL divergence (nats (e.g. primary 5vs12 and (reciprocal 12vs5)): 5/12dpf: 0.15 (0.135), 5/30dpf: 0.33 (0.261), 5/45dpf: 0.30 (0.247), 5/75dpf: 0.66 (0.539), 12/30dpf: 0.21 (0.13), 12/45dpf: 0.16 (0.099), 12/75dpf: 0.43 (0.26), 30/45dpf: 0.02 (0.022), 30/75dpf: 0.07 (0.074), 45/75dpf: 0.11 (0.072), and average y-depth (cm): 5dpf: 7.8, 12dpf: 7.1, 30dpf: 6.3, 45dpf: 6.4, 75dpf: 5.4, see also **Table 2** and **Table S1**). When quantifying the distribution of larval positioning over the first five minutes, the consequences of the behavioral transitions become evident: embryos and young planula (5 and 12dpf) first sink close to the bottom of the chamber before ascending. The homogeneity of the tracks at this age is striking, suggesting a passive rather than an active mode of dispersal. Generally, the first three ages have a trend towards the bottom, with sharper peaks for both bottom and top residence times at 30dpf as shown by depth distribution (**Fig. 2C,D**). Width distribution shows that most larvae are in the center of the arena (**Supplementary Fig. S1A** (bottom panel)) and track homogeneity during the first 30 seconds after insertion (**Supplementary Fig. S1D)** reveals the passive nature of a large proportion of the animals. This initial behavior is likely a startle response, common among invertebrate larvae in response to mechanical and light stimuli^21,22^.

In contrast, a clear shift in behavior towards active swimming is evident in older animals, exemplified through regular spiraling and disorganized trajectories. Swimming endows the animals, at least partially, with the option to choose in which direction to move. In our essay, this behavior seems to promote the exploration of the top (presumably a feeding driven behavior) or bottom (possibly akin to habitat choice) of the chamber, resulting in a more homogeneous width distribution (**Supplementary Fig. S1A**).

Is the position in the water column stable? Five minutes is a short timeframe, and we were curious about how the distribution develops over an extended period of time. Larvae were recorded for an additional 6 minutes (data not shown) and their position in the last minute (11th minute) plotted (**Fig. 2E**). This experiment corroborates that embryos (5dpf) are neutrally buoyant, as their position in the water column distributes uniformly in depth and width. A pattern consistent with neutral buoyancy is still visible in a proportion of 12dpf animals, although many larvae have switched to residency in the top layer at this age. The 30, 45 and 75dpf larvae show peaks at bottom and water surface, with uniform distribution along the X-axis. 45 and 75dpf animals perform “dives” close to the water surface that can be interpreted as akin to food searching behavior, a mode most pronounced at this later stage.

Our data show notable developmental changes and adaptations in behavior. Embryos and young larvae disperse passively and only start directed motility at later stages. Larvae seem to actively swim under still water conditions but not under turbulent currents, as direct insertion (mechanical stimulation through handling) induces movement suspension. This behavior is consistent with an energy preserving strategy.

### Motility patterns and behavioral modes explain vertical positioning

How can larvae control their vertical positioning? There are two possibilities: through buoyancy, which may require longer-term metabolic changes, or through flexible active swimming, which can occur more rapidly. To exclude buoyancy, we developed a second assay focused on horizontal mobility. Buoyancy is minimized by filming the larvae in a shallow water layer (0.28 cm, **Fig. 3A**), allowing us to observe motility patterns in the horizontal plane and describe detailed aspects of locomotion for larvae of the same age groups used in the vertical assay **Fig. 3A,B** after^23^).

**Figure 3.**
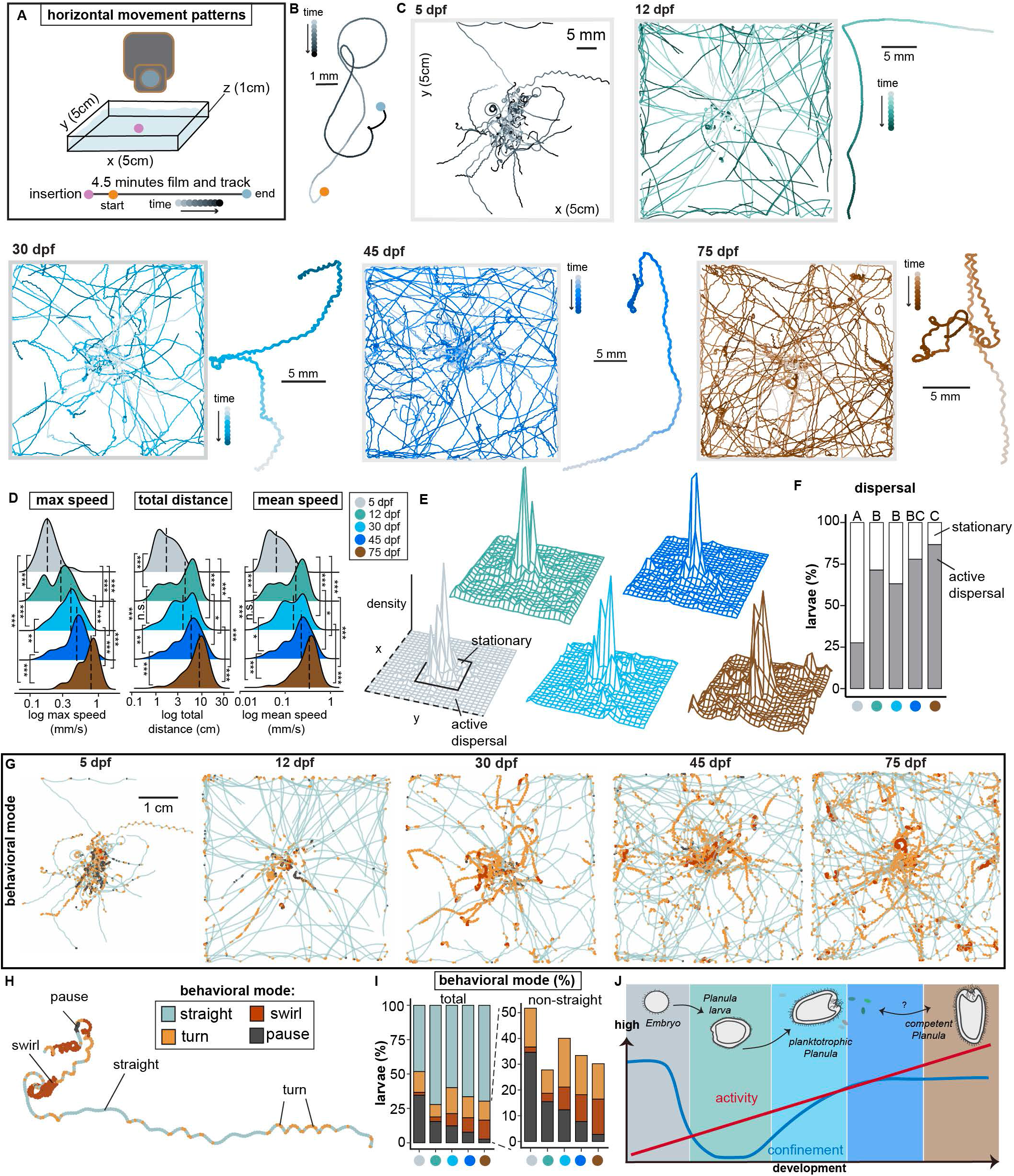
Motility state and behavioral mode define age dependent locomotion patterns. **A)** Experimental setup for horizontal tracking. **B)** Example track for horizontal behavior: insertion point (pink), experimental start point (orange) and end point (blue), color intensifies over the 4.5 minutes. **C**) Overlaid tracks of *Lophelia pertusa* larvae aged 5dpf (6N, 58n), 12dpf (6N, 63n), 30dpf (5N, 57n), 45dpf (6N, 68n), and 75dpf (5N, 53n), scale bar in the 5dpf arena applies to all arena plots. **D)** Distributions with median (dashed line): maximum speed (mm/s): 5dpf: 0.177, 12dpf: 0.28, 30dpf: 0.376, 45dpf: 0.468, 75dpf: 0.768), total distance (mm): 5dpf: 17, 12dpf: 43.5, 30dpf: 38.9, 45dpf: 58.8, 75dpf: 88.6), and mean speed (mm/s): 5dpf: 0.063, 12dpf: 0.169, 30dpf: 0.15, 45dpf: 0.224, 75dpf: 0.338). Statistical significance shown as ns p>0.05, *p<0.05, **p<0.001 and ***p<0.0001 by Kolmogorov-Smirnov test. **E)** Density plot per age group with area for “stationary” and “active dispersal” labeled as quantified in **F)** as average percent (Active dispersal (%): 5dpf: 27.59, 12dpf: 71.43, 30dpf: 63.16, 45dpf: 77.94, 75dpf: 86.79; stationary (%): 5dpf: 72.41, 12dpf: 28.57, 30dpf: 36.84; 45dpf: 22.06, 75dpf: 13.21). Different letters (A–C) indicate significant differences p<0.05 by two-tailed two-proportion z-test. See also **Table 3. G)** Tracks from **C** colored by behavioral state. **H)** Example track showing different behavioral modes which are quantified in percent in **I)** with all behavioral states (%) (left, straight: 5dpf: 48.20, 12dpf: 72.23, 30dpf: 59.93, 45dpf: 66.59, 75dpf: 69.81; turn: 5dpf: 15.14, 12dpf: 9.05, 30dpf: 18.90, 45dpf: 15.25, 75dpf: 13.76; swirl: 5dpf: 2.06, 12dpf: 3.28, 30dpf: 8.9, 45dpf: 10.5, 75dpf: 13.71; pause: 5dpf: 34.61, 12dpf: 15.44, 30dpf: 12.27, 45dpf: 7.67, 75dpf: 2.72). **J)** Summary of changes in larval behavior over developmental time. See also **Table 3**.

As suspected from the vertical assay, embryos (5dpf) and young planula larvae (12dpf) show relatively passive horizontal movements (**Fig. 3C,D** median mean speed (mm/s) 5dpf: 0.063, 12dpf: 0.169, median total distance (mm) 5dpf: 17, 12dpf: 43.5). 12dpf larvae show long-tailed (mean speed and total distance) or bimodal (maximum speed) distributions, suggesting that a major developmental transition takes place at this age. This is consistent with data from the vertical assay in which some larvae are still neutrally buoyant (similar to 5dpf), while others have assumed a more positively buoyant position (**Fig. 2E**).

On a global scale, larvae become faster, and travel further as they age (**Fig. 3C,D**: median maximum speed (mm/s): 5dpf: 0.177, 12dpf: 0.28, 30dpf: 0.376, 45dpf: 0.468, 75dpf: 0.768, median mean speed (mm/s): 5dpf: 0.063, 12dpf: 0.169, 30dpf: 0.150, 45dpf: 0.224, 75dpf: 0.338, median total distance (mm): 5dpf: 17.0, 12dpf: 43.5, 30dpf: 38.9, 45dpf: 58.8, 75dpf: 88.6). Metamorphosed animals showed a significant (p<0.0001, Mann Whitney test) reduction in mean speed relative to non-metamorphosed larvae of the same age (30dpf, **Supplementary Fig. S2A**, mean speed (mm/s) ±SEM larvae norm.: 0.167 ±0.013, metam.: 0.057 ±0.004), consistent with the movement seizure observed upon metamorphosis in other cnidarian larvae ^23^. This observation marks an important developmental bifurcation point with major implications for dispersal capacity: at one month (30dpf), animals are competent and can metamorphose, yet they can also maintain motility. Motile larvae between 1 and 2.5 months (30, 45 and 75dpf) show distinct swimming patterns compared to younger ages. At this more mature stage, animals swim with larger diameter spirals, with frequent directional changes, as clearly seen in the overlay and in individual tracks (**Fig. 3C**).

Larval activity-changes suggest that animals first disperse passively, most likely using currents, while later motility patterns are dominated by active swimming and more complex behaviors. To quantify the relevance of larval movement for active versus passive dispersal, we classified the larvae’s behavior as either “active dispersal” or “stationary” depending on whether or not the larvae left a perimeter threshold covering the inner 20% of the arena (**Fig. 3E**,**F)**. We found that active dispersal increases steadily over age, with the lowest proportion found at 5dpf and the highest proportion of animals leaving the inner perimeter at 75dpf (proportion active dispersal (%): 5dpf: 27.59, 12dpf: 71.43, 30dpf: 63.16, 45dpf: 77.94, 75dpf: 86.79, proportion stationary (%): 5dpf: 72.41, 12dpf: 28.57, 30dpf: 36.84, 45dpf: 22.06, 75dpf: 13.21; statistically significant ratios shown as compact letter display (A-C) for p<0.05 by two-proportion z-test, see also **Table 3**). These data strengthen the notion that larvae at certain developmental stages have a tendency towards movement suspension while others explore through active swimming.

Intricate behavioral transitions involving more sophisticated traits might not be captured by simple metric analysis. While maximum speed increased steadily with age, neither track mean speed nor total distance differ significantly between 12dpf and 30dpf (**Fig 3D**). To investigate if other aspects of swimming behavior change throughout larval development, we developed a more detailed analysis, distinguishing discrete behavioral modes and their location within the arena. Four distinct behavioral modes (“straight”, “turn”, “swirl” and “pause”, **Fig. 3G-I** and see material and methods) were identified and revealed that the larvae do not simply change speed. Larvae generally often swim linearly (**Fig. 3I**: average straight (%): 5dpf: 48.2, 12dpf: 72.23, 30dpf: 59.93, 45dpf: 66.59, 75dpf: 69.81), with a peak at 12dpf (**Supplementary Fig. S2C)**. When quantifying the time spent displaying other behavioral modes, we saw two clear trends: larvae “pause” less often over the course of development (average pause (%): 5dpf: 34.61, 12dpf: 15.44, 30dpf: 12.27, 45dpf: 7.67, 75dpf: 2.72) and increasingly started to “swirl”, a locally confined behavior that we interpret as search behavior (average swirl (%): 5dpf: 2.06, 12dpf: 3.28, 30dpf: 8.9, 45dpf: 10.5, 75dpf: 13.71). Turning behavior is relatively stable (average turn (%): 5dpf: 15.14, 12dpf: 9.05, 30dpf: 18.9, 45dpf: 15.25, 75dpf: 13.76) throughout development.

These observations lead to a model characterized by global developmentally increasing speed, but mode switching behavior (**Fig. 3J**). Larvae display early and late preferences in behavioral mode. Early movement is slower, confined, with more pauses, leading to passive travel through neutral buoyancy. Later movement is faster but locally confined due to frequent reorientations via swirls. This latter mode of behavior tends to occur when larvae are concentrated at the top or bottom of the water column (**Fig. 2E**) and may recapitulate a natural form of search behavior for food or settlement location.

### *Lophelia pertusa* larval anatomy changes during development

Another way of regulating positioning in the water column is by coordinating physiological transformations with active swimming modes to regulate behavior. To look for evidence of such coordination, we investigated developmental changes in subcellular architecture of *Lophelia pertusa* planulae by combining whole mount immunohistochemistry, toluidine blue-stained thin sections, and TEM imaging of larvae at three different ages: 12, 30, and 75 days post fertilization (dpf).

When examined externally (**Fig. 4A**), *Lophelia pertusa* larvae become increasingly elongated and densely ciliated throughout development, consistent with their ability as avid swimmers. Indeed, we observed an increasing cell density over time (**Fig. 4B,G,L**). The cilia have a typical 9+2 microtubule structure, both in the outer epithelium and in the oral cavity (**Fig. 4D-F**) and are surrounded by a crown of 11-13 microvilli (**Fig.4C-E, H** and **N)**, consistent with previous reports^13,15^.

**Figure 4.**
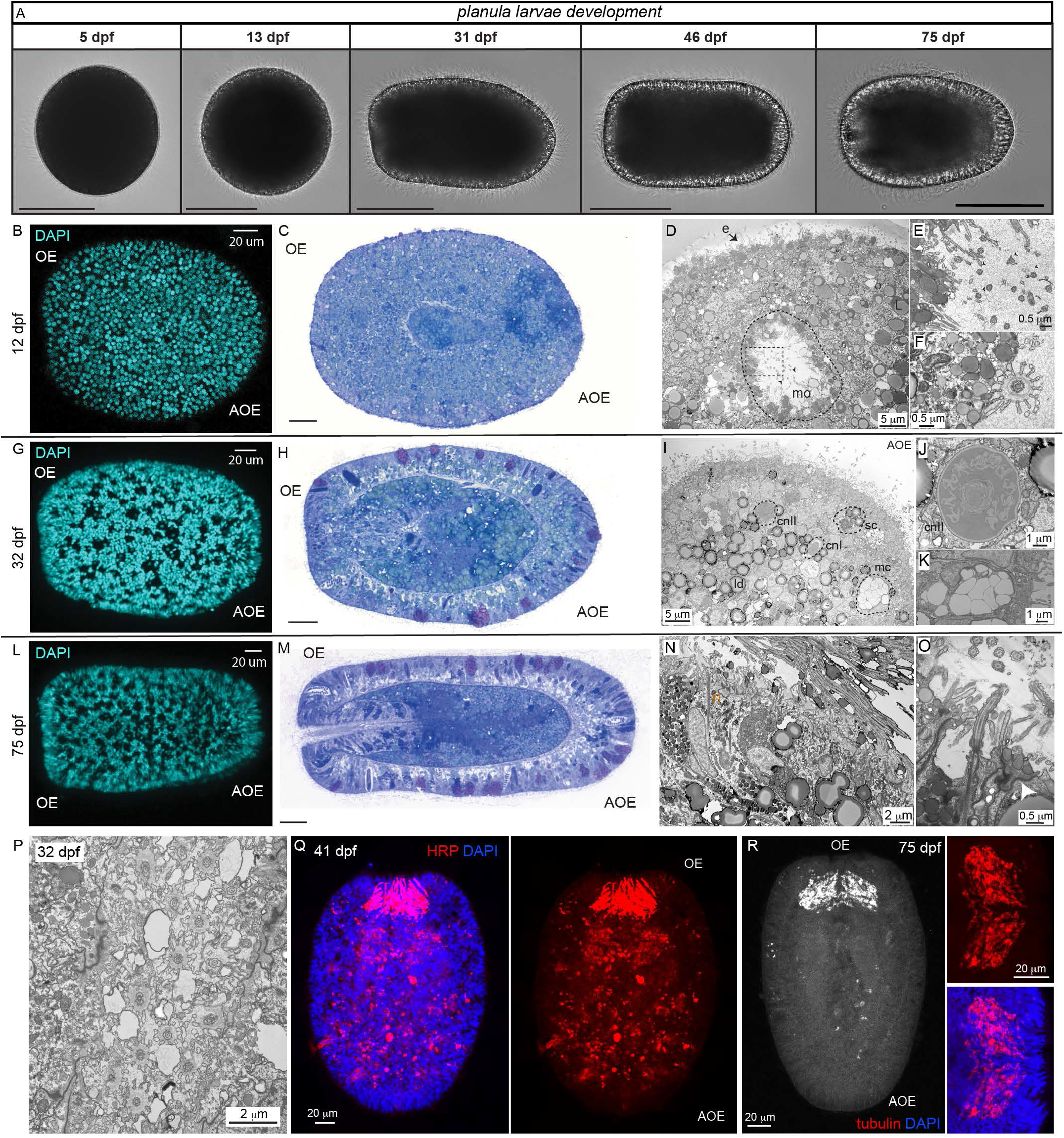
Larval anatomy changes over the course of development. **A)** Brightfield pictures of developing embryo and planula larvae at five different time points: 5, 12, 31, 46 and 75 days post fertilization (dpf). Scale bars: 100 μm. **B)** 12dpf old larvae stained with DAPI or **C)** section stained with toluidine blue showing lipid rich larvae (light green= lipid droplets). **D)** Lipid rich epithelium (e) and mouth opening (mo) at the oral end of the larva is clearly visible with 9+2 cilia also seen in inlet **E)** and **F)** magnification on base of motile cilium surrounded by crown of 13 microvilli. **G)** DAPI staining of 32dpf larvae showing larger number of cells and holes where large secretory cells are localized (basal nucleus visible in the middle). **H)** Toluidine blue staining showing a cross section with clear distinction into ecto- and endoderm, separated by the mesoglea. Distinct cell types can be clearly identified also seen in **I)** showing cnidocytes (cn) and secretory cells (sc) **J)** close-up of cnidocyte and **K)** secretory cell. **L)** 75dpf DAPI staining revealing more numerous secretory cells and elongation of the animal **M)** as also seen in the cross section. Extensive ciliation in the mouth opening and the reduced lipids can be clearly identified along with more numerous cnidocytes in the ectoderm and secretory cells in the endoderm (dark blue) which are magnified in **N)** where the epithelium with numerous and extremely long cilia is clearly visible interspaced with neurons (n), **O)** ciliated secretory cell and epithelium with potential gap junctions (white arrow). **P)** Cross section through cilia in oral cavity. **Q)** Stain with HRP reveals accumulation of sensory cells in mouth area as also confirmed in **R)** with tubulin staining. Scale bars: B,C,G,H,L,M= 20mm, OE= oral end, AOE= aboral end.

*Lophelia pertusa* larvae possess large deposits of lipids throughout the body (**Fig. 4** and **Supplementary Fig. S3A-C**), which appear to be metabolized throughout the 2.5 months of lifespan examined here (**Fig. 4C,H,M** (lipids in green) and corresponding EM **Supplementary Fig. S3A-C**). Compared to the 12dpf larvae, lipids in the 30 day old larvae have largely retreated into the gastric cavity and are even more sparse in 75 day old animals. Considering that vertical positioning in the water column is important for dispersal and feeding, the lipid content is likely to play a role in buoyancy control of the animals. Lipids are commonly used as a mechanism to regulate buoyancy in deep sea animals^24,25^ and the diminishing content of lipids observed in *Lophelia* over the 2.5 months is consistent with changes in water column positioning (**Fig. 2E**).

Due to the high lipid content, we were not able to perform histology before 12dpf. At this age, however, the larvae are elongated and full of lipids (**Fig 4B,C** and **Fig. S3A**), with no apparent differentiated cell types apart from ciliated epidermal cells. In contrast, 32dpf larvae show clear organization into ecto- and endoderm (gastroderm) (**Fig. 4H**), separated by the mesoglea (mg) (**Fig. 4H** and **Supplementary Fig. S3B)**. Several cell types primarily found in the ectodermal layer can be clearly identified at this stage: different types of cnidocytes (**Fig. 4I,J**) and secretory cells are present in high density, some with dense vesicles and some with large translucent chambers (**Fig. 4K**). Large secretory cells are easily identifiable by the “gaps” in the DAPI staining with a basal nucleus (**Fig. 4G**), and as purple stained cells in the toluidine sections (**Fig. 4H**).

Later in development (75dpf), some anatomical features of the larvae become more pronounced, especially the mouth opening and the pharynx region are densely ciliated and rich in secretory cells (**Fig. 4L** and **4M-O)**. The investment of *Lophelia pertusa* larvae in the feeding apparatus is evident through multiple lines of evidence: dense ciliation in the mouth area (**Fig. 4P**) and numerous secretory cells in the oral area. When using neuronal markers for invertebrate larvae, such as HRP (**Fig. 4Q**) and tubulin (**Fig. 4R**), a prominent “mouth brain” can be readily identified in the oral region. This nerve plexus has been previously described^13^ and underlines the investment of the planktotrophic larvae to sense suitable food sources. Furthermore, at 75dpf the larvae possess many different types of cnidocytes (**Fig. 4M** and **Supplementary Fig. S3C-E**), hinting towards an active predatory lifestyle, defense mechanisms, or readiness to anchor and attach to a specific substrate.

*Lophelia pertusa* larvae thus undergo massive anatomical transitions throughout their development. Changes in lipid content and appearance of cellular specializations are likely underlying some of the major behavioral transitions of the coral planula.

### Lifestyle and habitat determine larval behavior

Animals are well adapted to their specific ecological niche, however, whether larval behavior is adapted to dispersal mode is currently unknown. *Lophelia pertusa* reefs are found worldwide, many with significant connectivity^12^ supported by long-lived, planktotrophic and highly dispersive planula. To contrast this background with that of a species with very low larval dispersal capacity, we used larvae of the brackish anthozoan anemone *Nematostella vectensis*. The short-lived, lecithotrophic larvae of *Nematostella* encounter a closed niche in its estuarine habitats, preventing population connectivity geographically and making larval dispersal largely dispensable^26,27^ (**Fig. 5A-C**).

**Figure 5.**
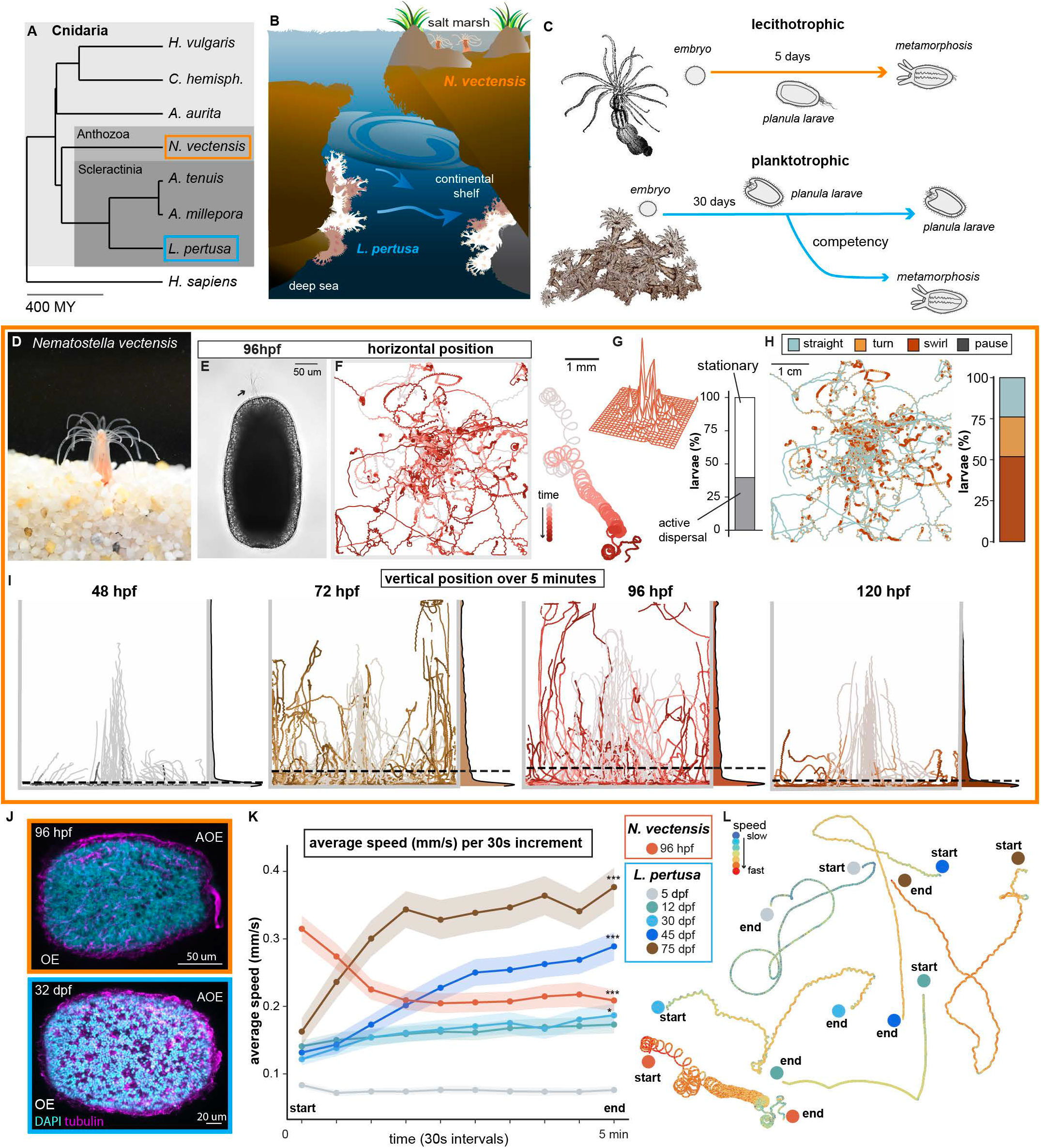
Dispersal behavior is shaped by lifestyle and habitat. **A)** Phylogenetic position of *Lophelia pertusa* and *Nematostella vectensis*. **B)** *Lophelia* is found in connected habitats of the deep sea on continental shelves, while *Nematostella vectensis* inhabits shallow pools in salt marshes which are geographically isolated. **C)** *Nematostella’s* lecithotrophic, poorly dispersing planula, metamorphose fast, as opposed to the slowly developing, planktotrophic, far dispersing *Lophlia* planula larvae. **D)** Adult *Nematostella vectensis* **E)** Brightfield image of a 96hpf larva, arrow= apical tuft (scale bar 50 µm). **F)** Overlaid tracks (9N,106n) and example track of 96hpf larvae. **G)** 3D density distribution and quantification of dispersal (active dispersal: 39.62%, stationary: 60.38%). **H)** Tracks colored by behavioral state. Percent (%) in each behavioral state: straight: 23.93, turn: 24.21, swirl: 51.84, pause: 0.02, scale bar 1cm. **I)** *Nematostella* larvae are mostly demersal at all ages as shown by overlaid tracks and density distribution of vertical position of 48hpf (26N, 111n), 72hpf (29N, 144n), 96hpf (28N, 142n) and 120hpf (25N, 137n). Density plots, with dotted lines at the mean (Mean depth (cm): 48hpf: 9.83, 72hpf: 9.19, 96hpf: 9.03, 120hpf: 9.76). **J**) Immunohistochemistry of a competent *Nematostella vectensis* (96hpf) and *Lophelia pertusa* (32dpf) larva stained with acetylated tubulin (magenta) and DAPI (cyan) (OE= oral end, AOE= aboral end). **K)** Mean speed over time shows opposing motor output to startle response (Mean speed (mm/s) ±SEM first and last timepoint *Nematostella vectensis* 96hpf: 0.312 ±0.02 and 0.209 ±0.016, p<0.001, Mann Whitney test), (Mean speed (mm/s) ±SEM at first and last timepoint *Lophelia pertusa* 5dpf: 0.083 ±0.004 and 0.076 ±0.006, p=0.064, 12dpf: 0.141 ±0.009 and 0.173 ±0.013, p=0.113, 30dpf: 0.122 ±0.009 and 0.187 ±0.017, p=0.024, 45dpf: 0.132 ±0.011 and 0.289 ±0.021, p<0.001, 75dpf: 0.16 ±0.02 and 0.377 ±0.028, p<0.001, by Mann Whitney test). Lines show the mean line ±SEM, see also **Table 5. L)** Individual example tracks colored by speed throughout 4,5 minutes of video. Start and end positions are labeled.

The nature of *Nematostella* (**Fig. 5D,E**) larvae’s poor dispersing capacity is evident immediately when analyzing behavior: horizontal swimming trajectories (**Fig. 5F-H**) show that swimming is highly confined, both globally (**Fig. 5G** and **Supplementary Fig. 4A**) and locally (**Fig. 5H**). The larvae behave 24.21% in turns and 51.84% in swirls, a local confinement behavior that might be interpreted as search behavior, and less than half of the animals (39.62%) leave the inner perimeter of the arena at all (**Fig. 5G**). When compared to the average swirl in *Lophelia* (average swirl between 2-13% and between 28-87% in “active dispersal” *Lophelia*), it becomes evident that these two ecologically distinct species employ opposite motility patterns.

*Nematostella* larvae show developmentally timed sensory perception, leading to pronounced dispersal behavior between 72-96hpf ^23^, a pattern that is matched in the vertical positioning (**Fig. 5I, Supplementary Fig. 4B-D** and **Table S4)**. To our surprise, when comparing the first 30 seconds of motility in the vertical plane, the trajectories of sensory perceptive *Nematostella* larvae looked strikingly different from those of *Lophelia* (**Supplementary Fig. 4E**, compare also **Supplementary Fig. 1D)**. Since both cnidarian planulae possess sensory structures at mid planula stage (**Fig. 4** and **5J**), we decided to analyze their horizontal behavior over time (**Fig. 5K,L**). Indeed, as anticipated from the first 30 seconds of the vertical assay, we found opposing responses in the two anthozoan larvae: *Nematostella* shows an “activated” startle response, while all but the youngest (5dpf) *Lophelia* larvae start out with similar speed despite stark differences in their swimming capabilities (**Fig. 3D**). *Lophelia* seems to freeze upon handling and transfer, and for older ages (30-, 45-, and 75dpf), larvae start swimming only after several seconds (**Fig. 5K**, mean speed (mm/s) at first and last timepoint: *Nematostella vectensis* 96hpf: 0.312 and 0.209, p<0.001; *Lophelia pertusa*: 5dpf: 0.083 and 0.076, p=0.064, 12dpf: 0.141 and 0.173, p=0.113, 30dpf: 0.122 and 0.187, p=0.024, 45dpf: 0.132 and 0.289, p<0.001, 75dpf: 0.16 and 0.377, p<0.001, by Mann Whitney test, see also **Table 5**). We therefore conclude that developmentally timed perception of sensory stimuli is conserved across these two anthozoans, but the behavioral output to these cues seems adapted to their natural niche. The same sensory input promotes passive dispersal under turbulent conditions for *Lophelia* but encourages active exploration for *Nematostella*.

## Discussion

Coral planula larvae provide an otherwise sessile species with the means to disperse and conquer new habitats, thereby maintaining its resilience and adaptability. Motility, however, is energetically costly^10^ and the acquisition and consumption of energy must be balanced to achieve an optimal trade-off between dispersal and retention. Movement of microscopic planulae can occur via different modes at different scales and many questions remain unanswered. Is the organism travelling passively, dispersing over kilometers, or actively localizing food or habitat on a mm-cm scale^8,9^? How are development, physiology, and behavior interacting to achieve “the right” dispersal-retention balance for the species?

Here, we provide a systematic description of developmental changes in anatomy and motility of the cold-water coral *Lophelia pertusa* planula. We find that the anatomical development supports increasingly complex motility patterns, which in turn influence the vertical positioning of the larvae, a critical determinant of dispersal. Sensory perception appears at later developmental stages, together with specialized effector cells. Developmentally timed sensory perception is conserved across the two anthozoan with naturally different dispersal behaviors that are examined here, but locomotor changes in response to stimuli are opposite, likely shaped to the habitat and dispersal style of these species.

Because the location of adult *Lophelia pertusa* colonies is dependent on food fall^28^ and *Lophelia* larvae consume food^14^, nutrient availability is likely to act as a strong settlement determinant. Feeding has been shown to alter morphology and is required for several planktotrophic invertebrate larvae to settle and metamorphose^29,30^ and self-generated flow increases nutrient uptake in planktotrophic larvae like *Lophelia*^10^. Our data support a large role for feeding, reflected anatomically by investment in the mouth opening, and behavioral drive to reach food rich planktonic layers. Coral larvae can survive for prolonged periods in the water column, but settlement success declines with time, because it is affected by nutritional state, specifically in planktotrophic planula^7,29,31^. Considering that nutrient uptake will enable prolonged residency in the water column and consequently have a major impact on the dispersal potential, this aspect is of major interest for future research.

One of the most energy consuming processes in the larvae is locomotion via motile cilia^10^. Mitochondria are commonly associated with cilia^32^ and their number scales with energy demand^33^. In *Lophelia* larvae, we find large accumulations of mitochondria in ciliated epidermal cells, underlining the energetic expenditure of active swimming. Some of these are peridroplet mitochondria (PDMs), organelles which may have a function in regulating the buoyancy of the animals, since PDMs can switch between lipid droplet accumulation and consumption depending on cellular state^34^. LDs are rich in triacyl glycerides, which directly regulate buoyancy, making this an exciting future avenue for investigating the feedback between metabolic regulation and motility^25^.

Reserving energy by passive travel is a simple way to conserve resources. Another way is by adapting sensory responses to sensory input. Coral larvae are often exposed to large water masses with strong currents in the open ocean. Suspending swimming activity upon mechanical stimulation and resuming active swimming only in still water conditions can be a natural way to balance energy availability and state-dependent needs of the species. In *Nematostella*, which generally experiences less flow-induced stimulation in its natural environment of secluded estuaries, we observed an opposite response. Consistent with our observations, recent studies have reported changes in coral swimming when exposed to ecologically relevant sensory information^8,22^. Developmentally timed sensory structures promote stage-specific behavior in marine invertebrate larvae thereby serving as another mechanism to restrain energy expenditure ^23,35–37^. The interplay between metabolism and sensation would be a plausible mechanism to choose a suitable habitat for settlement in cold-water corals.

Our data show that the behavior and physiology of cnidarian planula larvae is more complex than previously appreciated. The comparative analysis between *Nematostella vectensis* and *Lophelia pertusa* revealed some habitat specific adaptations that are shaped by different “drives” over development in both species. Such developmentally timed, niche adapted behaviors might allow optimal use of energy resources and contribute to maximal dispersal success. Older larvae swim more actively, with a position in the water column changing from neutrally buoyant to exploratory via active swimming in the experimental arena. Underlying anatomic developmental changes seem consistent with the motility patterns: young animals are extremely lipid rich and neutrally buoyant. As the lipids are metabolized, animals become increasingly active and swim to their chosen position in the water column. Lipids can be selectively metabolized when needed, and different lipids have different effects on buoyancy^25^. The resulting consequences on vertical position could have major effects on the larvae’s dispersal and subsequent recruitment^19^. Dispersal models have assumed positively buoyant larvae^12^ and locomotion mode appears to have little weight in the model larvae^38^. Consistent with this, our results suggest that the dispersal phase in *Lophelia* planula is largely passive, reinforced by ciliary arrest upon mechanical stimulation. An active swimming phase, previously not considered in existing models, may serve food and settlement localization in relatively still waters. The characterization of this developmentally timed and sensory-driven behavior is a major step forward in our understanding of how microscopic larvae achieve reliable dispersal and conquest of novel habitats. Dynamic aspects of the larvae’s motility over the course of development should be considered in future dispersal models.

## Methods

### Animals

#### Lophelia Pertusa (Syn. Desmophyllum pertusum)

Coral individuals were sampled with an ROV and net from the Gunnerus at the Geitneset reef (between 150 and 300m depth, in 2022 and 2024). Animals were kept in a cold room (7°C) at Trondheim biological station (TBS) in 50 -100L tanks with a flow-through water system (water from 100m depth) and additional water pump (Aqua Medic SmartDrift3.1). Animals were fed with a mix of copepods and barnacle larvae (Planktonic As) every 2-4 weeks.

For spawning, individuals were cleaned, broken into smaller fragments, and mixed to ensure maximal fertilization success. Spawning adults from two seasons were used in this study (Spring 2023 and 2025). Fertilized eggs were collected from the tanks and transferred to glass jars where they were maintained and washed regularly with filtered seawater (FSW).

#### Nematostella vectensis

Animals were maintained as previously described ^23^: in brief the sea anemones were kept in glass Pyrex dishes filled with 14 ppt ASW (Red Sea Salt, Red Sea Aquarium System) at 18°C in the dark. The anemones were induced to spawn regularly and after fertilization and development at RT (21°C), larvae and polyps were used at indicated ages for specific experiments.

### Behavior

Unless stated otherwise, all behavioral experiments were filmed using a Nikon Z50DX16-50 camera with a Nikkor MC105/2.8S lens at 24fps. To illuminate the chambers, several 8-RGB LED NeoPixel Sticks (Adafruit, product ID1426) were used, and controlled through an Arduino Mega2560 Rev3 and Arduino IDE2.2.1 software, resulting in 180lux (in air) by wall edge and 26 lux (in air) in the middle for vertical chamber (RGB setting 2,2,2 and no diffuser paper) and 150lux (in air) by edge wall and 69lux in the middle (in air) in horizontal chamber (RGB setting 3,3,3 with diffuser paper).

#### Horizontal swimming

Experiments were performed in accordance with^23^ with *Nematostella* (96 hours post fertilization (hpf)) at RT, and *Lophelia* larvae (5-, 12-, 30-, 45- and 75- days post fertilization (dpf)) in a cold room set to 7°C. Approximately 12 larvae were transferred into the center of a custom-made chamber (1×5×5cm) containing 5mL of 14ppt ASW at RT for *Nematostella* and 7mL of 0.2μm filtered sea water (FSW) at 7°C for *Lophelia*, to mimic the natural conditions for each species. Videos of 5dpf and 12dpf Lophelia larvae were recorded using a GoPro Hero10. Larvae were checked under a microscope before each experiment. In the metamorphosed group almost all larvae showed signs of developing tentacle buds.

#### *Horizontal behavioral analysis* (Particle tracking)

Videos were analyzed according to^23^. Individuals that collided and could not be reliably distinguished were excluded. Gaps were interpolated by linking the coordinates flanking a gap when possible. Because movement (e.g. mean speed) changed over time and fragmented tracks could bias summary metrics, only near-complete tracks (>88% of video length; ≥5760/6480 frames; 95.06% of all detected tracks) were used. Gaps larger than 168 frames were excluded from calculations of mean speed, total distance travelled, and maximum speed to reduce artifacts from intermittent detection loss. A brief camera displacement during the first ∼30 seconds after larval introduction caused a temporary change in coordinates. Consequently, data were analyzed in two segments (5-30s and 30-300s). Mean speed for 5-30 seconds was calculated separately.

We also defined maximum speed as the highest average velocity over a 10 frame (0.416s) rolling window within each track, reducing the effect of tracking errors. (I) Maximum speed= max_i_ (1N_i_ ^−1^ • Σ (d_j•j + 1_ • s^-1^)) Where (d_j•j + 1_ • s^-1^) is the momentary speed at index j, W is the window length, and N_i_ is the number of samples in the window. Path straightness was assessed using the confinement ratio, defined as the net displacement divided by total distance. To accurately capture path straightness, the ratio was calculated within consecutive 30s intervals, and then averaged to yield a final confinement ratio per track. Data were analyzed in GraphPad Prism for Windows (GraphPad software) and R Version 4.3.1 (https://www.R-project.org/).

#### Vertical swimming

*S*wimming behavior was performed with *Nematostella* larvae (48-, 72-, 96- and polyps at 120hpf) at RT, and *Lophelia* embryos and larvae (5-, 12-, 30-, 45- and 75dpf) in a cold room set to 7°C. In each experiment, approximately 6 larvae were carefully transferred to the center of a custom-made chamber (100×150×10mm), filled with either 100mL of RT *Nematostella* media or 7°C *Lophelia* media respectively. 5-minute (*Nematostella*) and 11-minutes videos (5+6min; *Lophelia*) were recorded. To account for telecentricity and perspective errors, the camera was placed 140cm away from the chamber.

#### Vertical behavioral analysis (Particle tracking)

Videos were tracked like previously described, and coordinate information was utilized for analysis. Tracks from individuals stuck to air bubbles were excluded. Due to the assay’s inferior contrast between larvae and background, fragmented tracks were not stitched together. Due to minor effects of telecentricity and resolution, the bottom of the chamber was found to be at 10 ±0.2cm depth, which was standardized to 10cm for the following analyses. The average depth per experiment and per age was calculated for 0-5 minutes for both species, as well as the 11^th^ minute for the *Lophelia* dataset.

### Histology

#### Immunohistochemistry

*Lophelia* larvae were fixed in 4% paraformaldehyde (PFA) in FSW for 16-24hr on a rocker at 7°C. Larvae were subsequently washed with PBST (TritonX 0.3%, 10-15 times) and incubated in primary AB (anti acetylated Tubulin 1:100 in PBST) at 4°C 12-48h. After subsequent washes PBST (10-15 times), antibody solution containing secondary ABs was applied overnight on a rocker at 4°C, samples were subsequently washed with PBST and DAPI (1:1000) was applied for 30 min at 4°C. Finally, the samples were washed 3-5 times in PBS, mounted in ProLong™ Glass Antifade Mountant (Thermo Fisher scientific, P36980), and imaged with a Zeiss LSM800 confocal. Images were processed using FIJI and Adobe Photoshop. *Nematostella* larvae were stained as previously described in^23^

#### EM

##### Primary fixation

Larvae of indicated ages were fixed in FSW with double concentration fixative 5% glutaraldehyde + 4% paraformaldehyde until the larvae sank to the bottom of the tube. Samples were centrifuged and the supernatant removed, 2,5%glutaraldehyde and 2%paraformaldehyde (in FSW) added to the pellet, fixed for 1-2 hours and then stored at 4°C.

##### Encapsulation in agarose

The samples were rinsed with FSW and encapsulated in 2% agarose in sea water. A razorblade was used to cut out the larvae into pieces. Excess agarose around the larvae was removed with a razorblade and the pieces were put in FSW.

##### For conventional TEM preparation

Post fixation was carried out for 1 hour with 2% osmiumtetroxide and 1,5% potassium ferrocyanide in FSW at RT, in the dark. Samples were rinsed with sea water and dehydrated with an ethanol series (VWR, 20821.310) 50%, 70%, 90% and 4 x 100% for 15 minutes at RT. Acetone (VWR, 83683.290) was used as a transitional solvent for (2×10 min RT), and subsequently infiltrated with an increasing concentration of epoxy LX-112 (Chemi-Teknik AS) in acetone; 1+2 for 1 hour, 1+1 for 1,5 hours, 2+1 for 2 hours. Infiltration with pure resin was carried out overnight (RT), samples were embedded in a mold with fresh resin and polymerized at 60°C for 3 days.

The blocks were trimmed using an ultramicrotome (Leica EM UC7) with a dry glass knife. Sections of 1µm were cut, put on a microscope slide and stained with toluidine blue for orientation using a light transmission microscopy. Ultrathin sections (60nm) were cut with an ultramicrotomy (Leica EM UC7) using a diamond knife (DiATOME) and subsequently collected on slot Copper grid with Formvar support film. Counterstaining was done with 4% Uranylacetate in 50% ethanol for 25 minutes, and 1% Lead Citrate in 0,2 M NaOH for 5 minutes.

##### For Serial block-face SEM sample preparation

The sample was stained *en block* using an established protocol for microwave (Biowave Pro+, Ted Pella). The procedure included successive exposures of the sample to reduced OsO4, thiocarbohydrazide (TCH), OsO4, uranyl acetate and lead aspartate. The samples were then dehydrated and infiltrated in epoxy resin. Finally the samples were embedded in silicone mold with fresh resin and polymerized at 60°C for 3 days. Blocks were trimmed using an ultramicrotome (Leica EM UC7) with a dry glass knife. Sections of 1µm were cut, put on a microscope slide and stained with toluidine blue for orientation using a light transmission microscopy. The area of interest was then selected for ultrathin sectioning. Ultrathin sections (60nm) were cut with an ultramicrotomy (Leica EM UC7) using a diamond knife (DiATOME) and subsequently collected on slot Copper grid with Formvar support film.

##### Imaging

Larvae of the desired age were selected at random and placed individually on a glass coverslip in 0.2μM filtered FSW for *Lophelia pertusa* or 0.2μM filtered 14ppt ASW for *Nematostella vectensis* in a light squish-prep to restrict larval movement in the z-plane. Larvae were then imaged with a 20x objective of a Nikon Eclipse Ti2-U inverted microscope fitted with a Hamamatsu Orca-flash 4.0 camera (model C13440-20CU).

## ACKNOWLEDGEMENTS

We thank the crew of the Gunnerus for sampling *Lophelia Pertusa* individuals and the IKOM EM facilities, specifically Thi My Linh Hoang for troubleshooting TEM in *Lophelia* larvae. This research was supported by the ERC StG “EnvIronchannel” (101076516) granted to L.v.G.

## AUTHOR CONTRIBUTIONS

M.L and J.H performed behavioral experiments and analysis, M.M.S performed imaging experiments, L.v.G and M.L performed immunohistochemistry, L.v.G acquired funding. All authors helped with animal maintenance, conceptualized the study, analyzed the data, designed and created the figures, and wrote the manuscript.

## Supplementary Figures

**Supplementary Figure 1.**
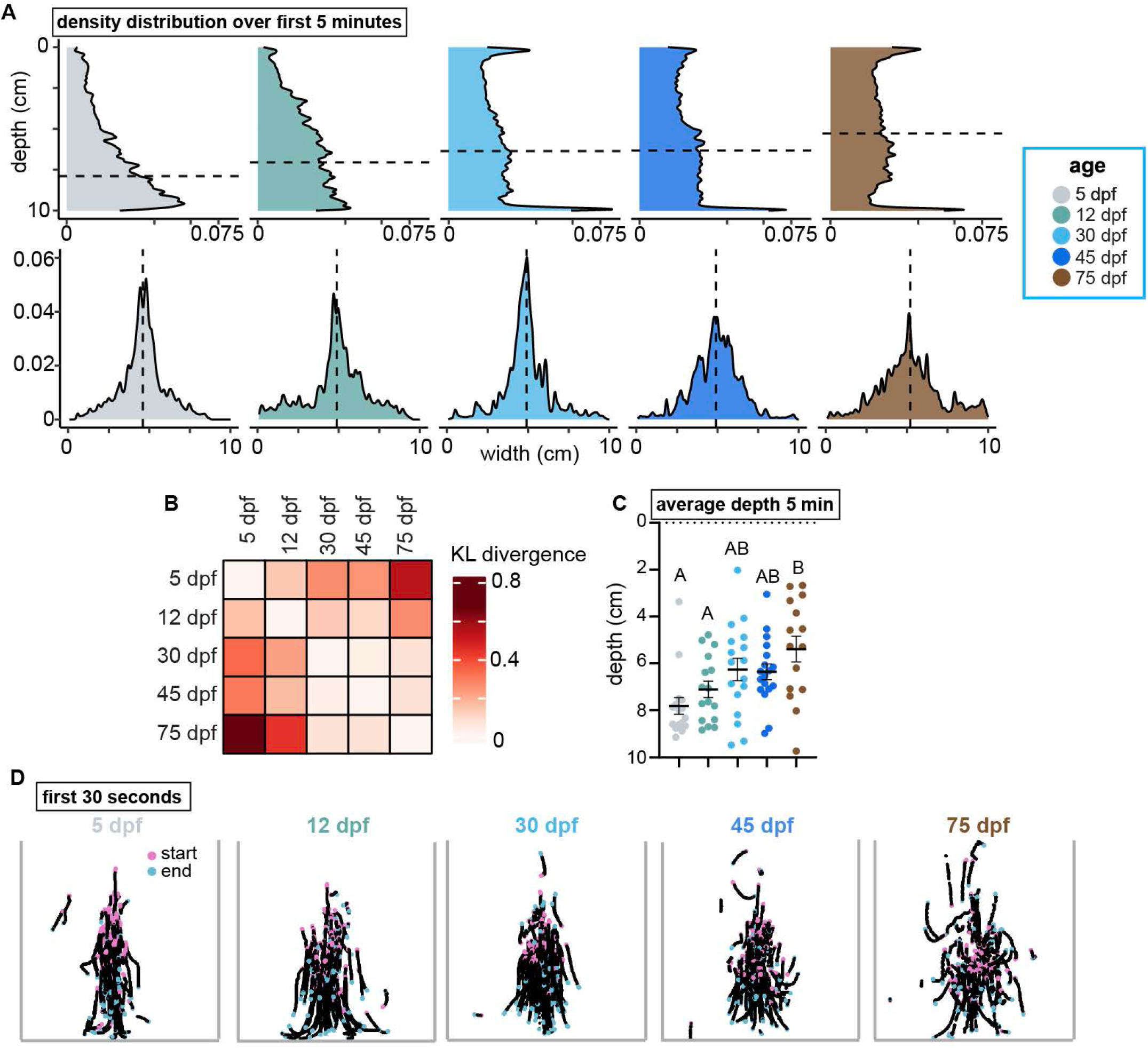
Details of vertical distribution of *Lophelia pertusa* planulae. **A)** Density plots show the depth (x-axis square root transformed) and width distribution during the first five minutes. Dotted lines at mean (Mean depth (cm): 5dpf: 7.89, 12dpf: 7.04, 30dpf: 6.36, 45dpf: 6.33, 75dpf: 5.28; mean width (cm): 5dpf: 4.60, 12dpf: 4.88, 30dpf: 4.90, 45dpf: 4.90, 75dpf: 5.21). **B)** Matrix of Kullback-Leibler tests shows the KL-divergence (nats) between vertical distributions of different ages for 0-5 minutes (see also **Table S1**). **C)** 0-5 minute average depth (cm) calculated per experiment, lines at mean ±SEM (5dpf: 7.81 ±0.36, 12dpf: 7.10 ±0.35, 30dpf: 6.25 ±0.48, 45dpf: 6.35 ±0.33, 75dpf: 5.38 ±0.55). Statistical significance of p<0.05 is denoted by different letters by ANOVA and Tukey’s multiple comparisons test. **D**) A segment of the first 30 seconds of all larval tracks for 5dpf (16N, 89n), 12dpf (16N, 99n), 30dpf (18N, 113n), 45dpf (17N,106n), and 75dpf (15N, 105n). Pink and blue spots indicate the 1st second and the 30th second, respectively.

**Supplementary Figure 2.**
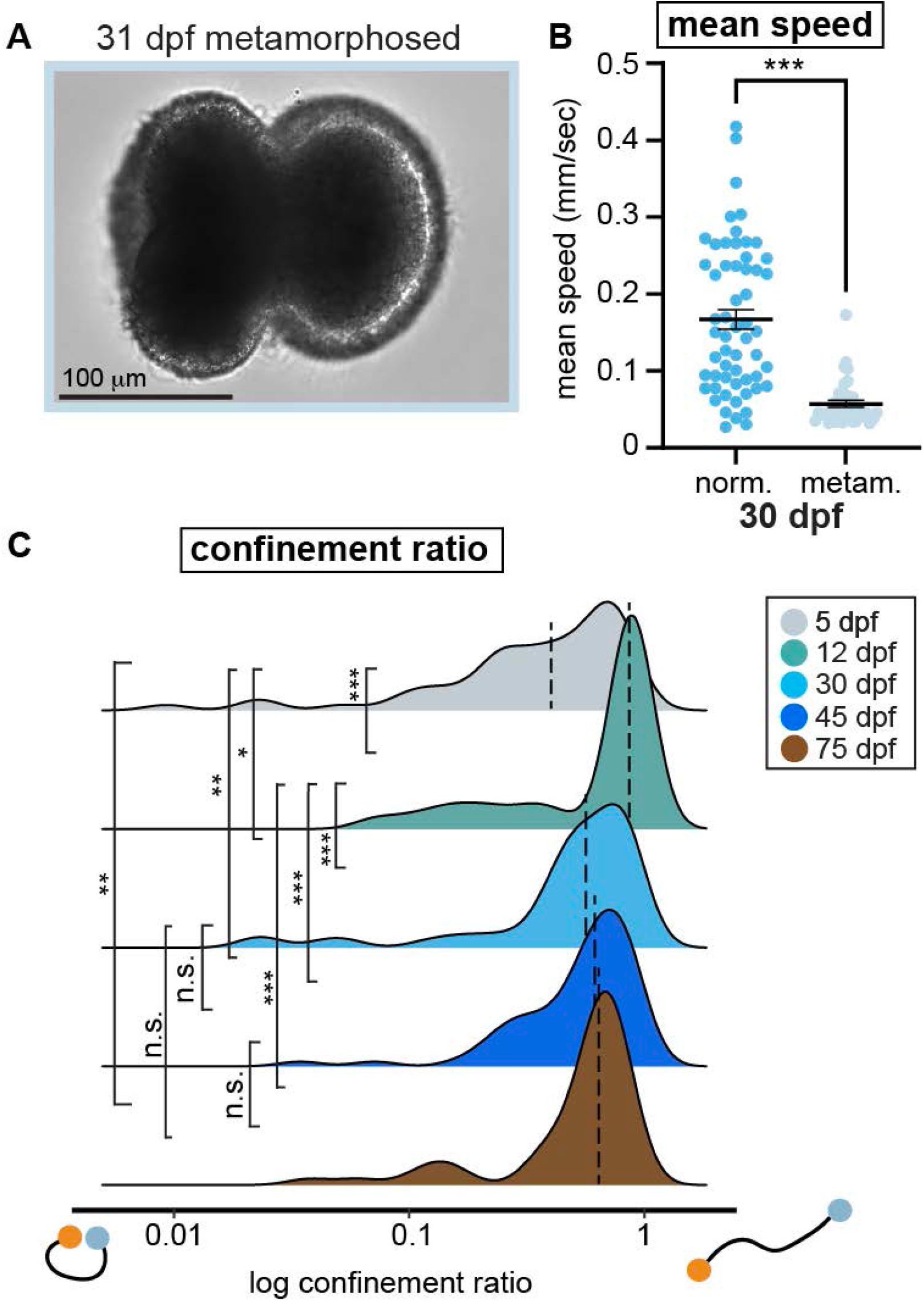
Behavior in metamorphosed animals and quantification of dispersal. **A)** Brightfield Image of a metamorphosed *L. pertusa* larva (31dpf). **B)** Mean speed of normal (norm.) planula larvae (5N, 57n) and metamorphosed larvae (metam.) (4N, 39n) from 4,5 min of video recording around one month of age lines at mean ±SEM (norm.: 0.167 ±0.013, metam.: 0.057 ±0.004. Statistical significance (p<0.001) denoted by asterix using Mann-Whitney test. **C)** Distributions of confinement ratio (dotted line at median): 5dpf (6N, 58n): 0.402, 12dpf (6N, 63n): 0.861, 30dpf (5N, 57n): 0.564, 45dpf (6N, 68n): 0.615, 75dpf (5N, 53n): 0.638. Statistical significance shown as ns p>0.05, *p<0.05, **p<0.001 and ***p<0.0001 by Kolmogorov-Smirnov test. See also **Table S2**.

**Supplementary Figure 3.**
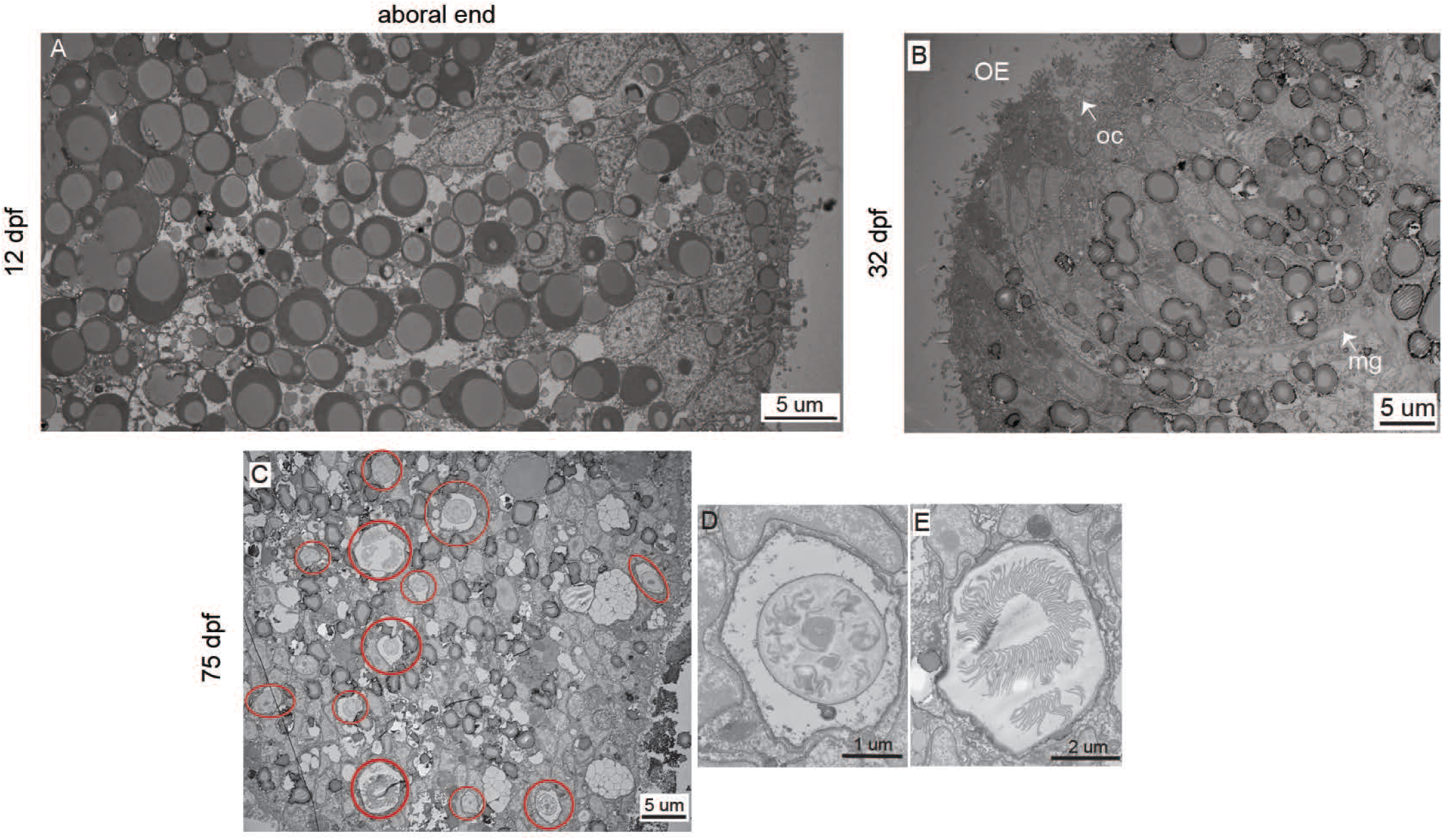
Anatomical details of EM sections in *Lophelia pertusa* larvae. **A)** The aboral end shows prominent lipid droplets distributed over the surface of the larvae (12dpf). **B)** Oral end (OE) of 32dpf old larvae with markedly reduced lipid content. Mesoglea (mg) and different cell types are readily identified at this age. Oral cilia (oc) mark the opening of the gastric cavity. **C)** At 75 days many different cnidocytes can be observed (red circles). **D)** and **E)** close up of cnidocytes.

**Supplementary Figure 4.**
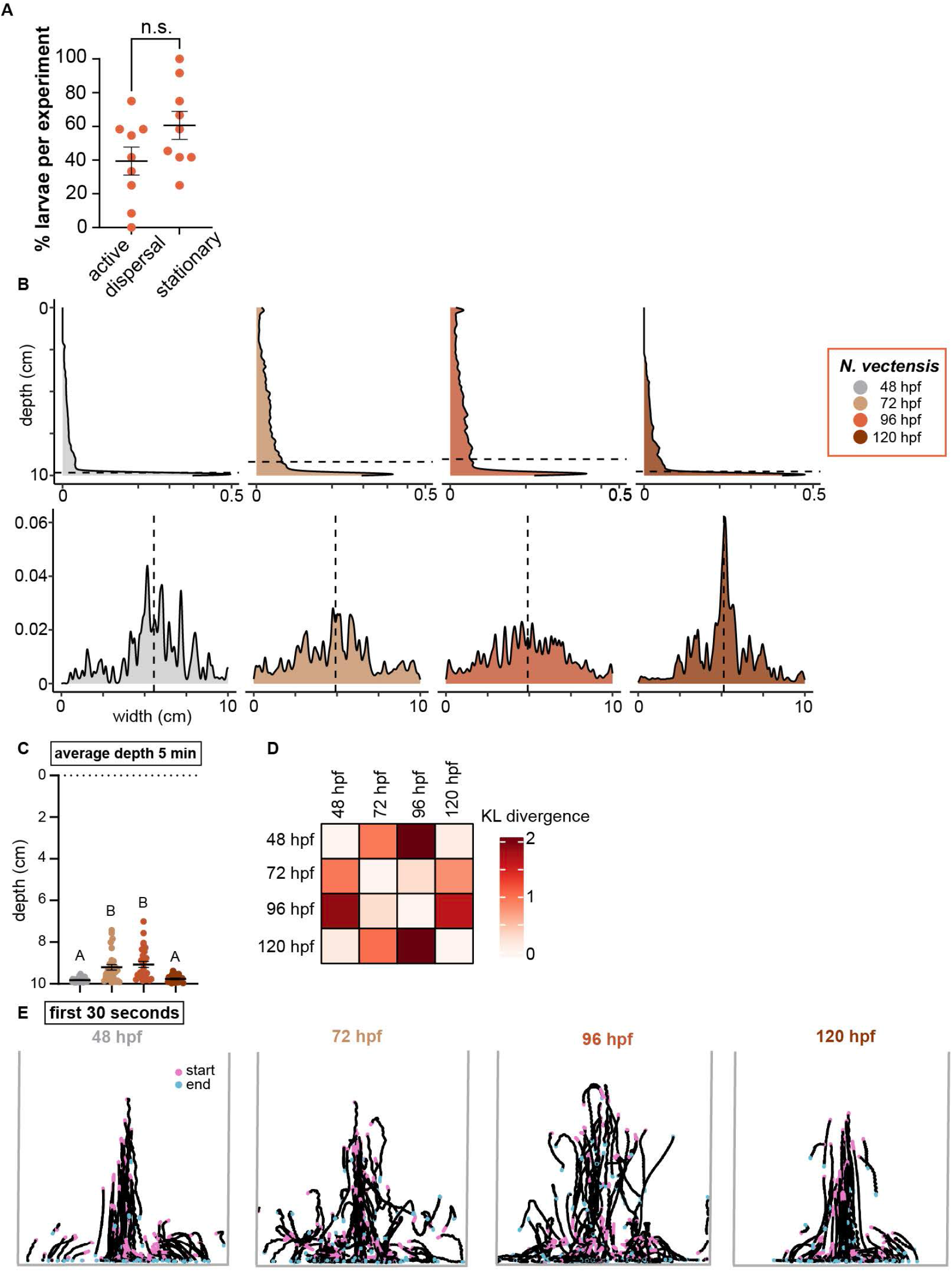
Quantification for *N. vectensis* behavior. **A)** Quantification percentage (per experiment, 9N) of 96hpf *Nematostella vectensis*, lines at mean ±SEM classified into active dispersal (39.39% ±8.3) or stationary (60.61% ±8.3). Significance (p=0.238, paired t-test) **B)** Density plots showing the average depth-(x-axis is square root transformed) and width-distribution the first five minutes after insertion for all larvae at 48hpf (26N, 111n), 72hpf (29N, 144n), 96hpf (28N, 142n) and 120hpf (25N,137n). Dotted lines show the mean depth and width (Mean depth (cm): 48hpf: 9.83, 72hpf: 9.19, 96hpf: 9.03, 120hpf: 9.76; mean width (cm): 48hpf: 5.55, 72hpf: 4.92, 96hpf: 4.91, 120hpf: 5.14). **C)** Average depth calculated per experiment, lines at mean ±SEM. Mean depth (cm): 48hpf: 9.82 ±0.021, 72hpf: 9.21 ±0.13, 96hpf: 9.07 ±0.145, 120hpf: 9.76 ±0.034). Statistical significance of p<0.05 denoted by different letters by Kruskal-Wallis test and Dunn’s multiple comparisons test. **D)** Matrix of Kullback-Leibler tests shows the KL-divergence (nats) between vertical distributions of different ages during 0-5 minutes: (e.g. primary 48vs72 and (reciprocal 72vs48)): 48/72hpf: 0.95 (0.95), 48/96hpf: 2.08 (1.89), 48/120hpf: 0.09 (0.11), 72/96hpf: 0.28 (0.27), 72/120hpf: 0.79 (1.02), 96/120hpf: 1.65 (2.06). **E)** A segment of the first 30 seconds of all larval tracks for 48hpf (26N, 111n), 72hpf (29N, 144n), 96hpf (28N, 142n) and 120hpf (25N,137n). Pink and blue spots indicate the 1st and 30th second, respectively. See also **Table S4**.

